# Spatiotemporal dynamics of zygotic genome activation in basal chordates revealed by interspecific hybrids

**DOI:** 10.1101/2022.06.01.494324

**Authors:** Jiankai Wei, Wei Zhang, Liang Leng, An Jiang, Yuting Li, Yonghang Ge, Quanyong Zhang, Liya Zhang, Haiyan Yu, Kai Chen, Bo Dong

**Affiliations:** Sars-Fang Centre, MoE Key Laboratory of Marine Genetics and Breeding, College of Marine Life Sciences, Ocean University of China, Qingdao 266003, China; Laboratory for Marine Biology and Biotechnology, Qingdao National Laboratory for Marine Science and Technology, Qingdao 266237, China; Chengdu University of Traditional Chinese Medicine, Chengdu 611137, China; State Key Laboratory of Primate Biomedical Research and Institute of Primate Translational Medicine, Kunming University of Science and Technology, Kunming, Yunnan 650500, China; Institute of Evolution & Marine Biodiversity, Ocean University of China, Qingdao 266003, China

**Author notes:** Correspondence, (K.C.); (B.D.).

**Keywords:** zygotic gene activation, ascidian, hybrids, transcriptome, single-cell RNA-sequencing

## Abstract

Zygotic genome activation (ZGA), a universal process in early embryogenesis that occurs during the maternal-to-zygotic transition, involves reprogramming in the zygotic nucleus that initiates global transcription. In recent decades, knowledge of this process has been acquired from research on various model organisms; however, a consensus explanation of the mechanism underlying the process, especially in relation to housekeeping gene reactivation, is lacking. Here, we used hybrids derived from two ascidian species (*Ciona robusta* and *C. savignyi*), which diverged >120 Mya with significant divergence among most orthologous genes, to symmetrically document the unique dynamics of ZGA in urochordates. We found two co-ordinated waves of ZGA, representing early developmental and housekeeping gene reactivation, during the 8-cell to 110-cell stage. Comparative analysis revealed the regulatory connection between maternal and zygotic genes as well as allelic-specific expression in a species-rather than parental-related manner, which was attributed to the divergence of cis-regulatory elements. Single-cell RNA sequencing revealed that spatial differential reactivation of paternal housekeeping genes was significantly correlated with the mechanical property of each cell type. These findings potentially provide a new system for understanding the evolution and adaptation of strategies regulating ZGA in basal chordates.

## Introduction

Despite divergent cleavage patterns among metazoans, the general molecular process of initializing developmental programming is highly conserved with few exceptions, even in plants. Following fertilization, the zygotic genome is initially quiescent, but some developmental genes are transcribed after a few rounds of division during cleavage, and widespread zygotic genome activation (ZGA) occurs coincident with housekeeping gene reactivation. Stepwise ZGA and transient silencing of housekeeping genes have been documented in several model species, suggesting that this process has been preserved under extreme selection pressure and plays important role in embryogenesis^1^. Knowledge of this process has largely been acquired from studies of *Drosophila, Danio rerio*, and *Xenopus*, which show mid-blastula transition (MBT) during the massive ZGA that co-occurs with the transition from synchronous division to prolonged asynchronous division^2^. In an attempt to explain the mechanisms of ZGA, several models have been proposed, including the nuclear-to-cytoplasmic (N/C) ratio model^3^, activator-accumulation model^4^, and chromatin state dynamics model^5, 6^. However, ZGA process are distinct according to different developmental paradigms^7^, and the regulation of ZGA in animals that do not show MBT remains poorly understood. Therefore, research on ZGA in a range of organisms, especially those considered to lie on pivotal transition points during animal evolution, would provide new insights into the evolution and adaptation of the molecular strategies underlying the ZGA process.

Ascidians, one of the closest living relatives of vertebrates, have been used in embryonic research for more than a century, and the initiation of the developmental programming in ascidian embryos has been intensively studied^8-11^. Due to the strict mosaic embryogenesis, cell lineages of multiple ascidian species have been well characterized, and the genomes of these species have also been sequenced^12-15^. Among the ascidians studied, two co-occurring solitary species from genus, *Ciona robusta* and *C. savignyi*, have been widely used in gene regulatory network studies as models of embryogenesis and chordate evolution. Despite their conserved embryogenesis patterning, these species actually diverged around 122 (± 33) Mya^16^, which is similar to the divergence time of humans and kangaroos^16, 17^. Hybrids of *C. robusta* and *C. savignyi* have not been reported in nature because of the chorion membrane that prevents cross-species fertilization and self-fertilization^18^. Nevertheless, by enzymically removing the chorion membrane of eggs, hybrid embryos can be derived in both directions^18, 19^. To the best of our knowledge, these are the most distinct hybrid embryos obtained in multicellular organisms. Moreover, they provide the opportunity to study ZGA dynamics using natural “genetic tags,” which overcome the challenge of distinguishing zygotic transcripts from maternally deposited RNAs^20^.

In the present study, we use hybrids produced from cross-fertilization between *C. robusta* and *C. savignyi* as models to study the co-ordination and preference between maternal and paternal transcripts during ZGA. We also assessed the reactivation of housekeeping genes in different lineages during embryogenesis at a single-cell resolution using single-cell RNA-sequencing. Our results improve our understanding of the dynamics of ZGA and provide new insights into the regulatory mechanisms of this critical process.

## Results and discussion

Both *C. robusta* and *C. savignyi* are tunicate species with similar morphologies that are broadly distributed across most coasts around the world (Fig. 1a) but diverged ∼122 Mya (Fig. 1b), which is larger than the distance of any two extant placentals^16^. The interfertility of these two highly divergent species was reported in a previous study^21^. Therefore, the hybrid from those *Ciona* species provides an excellent model with which to study hybrid embryos.

**Fig. 1.**
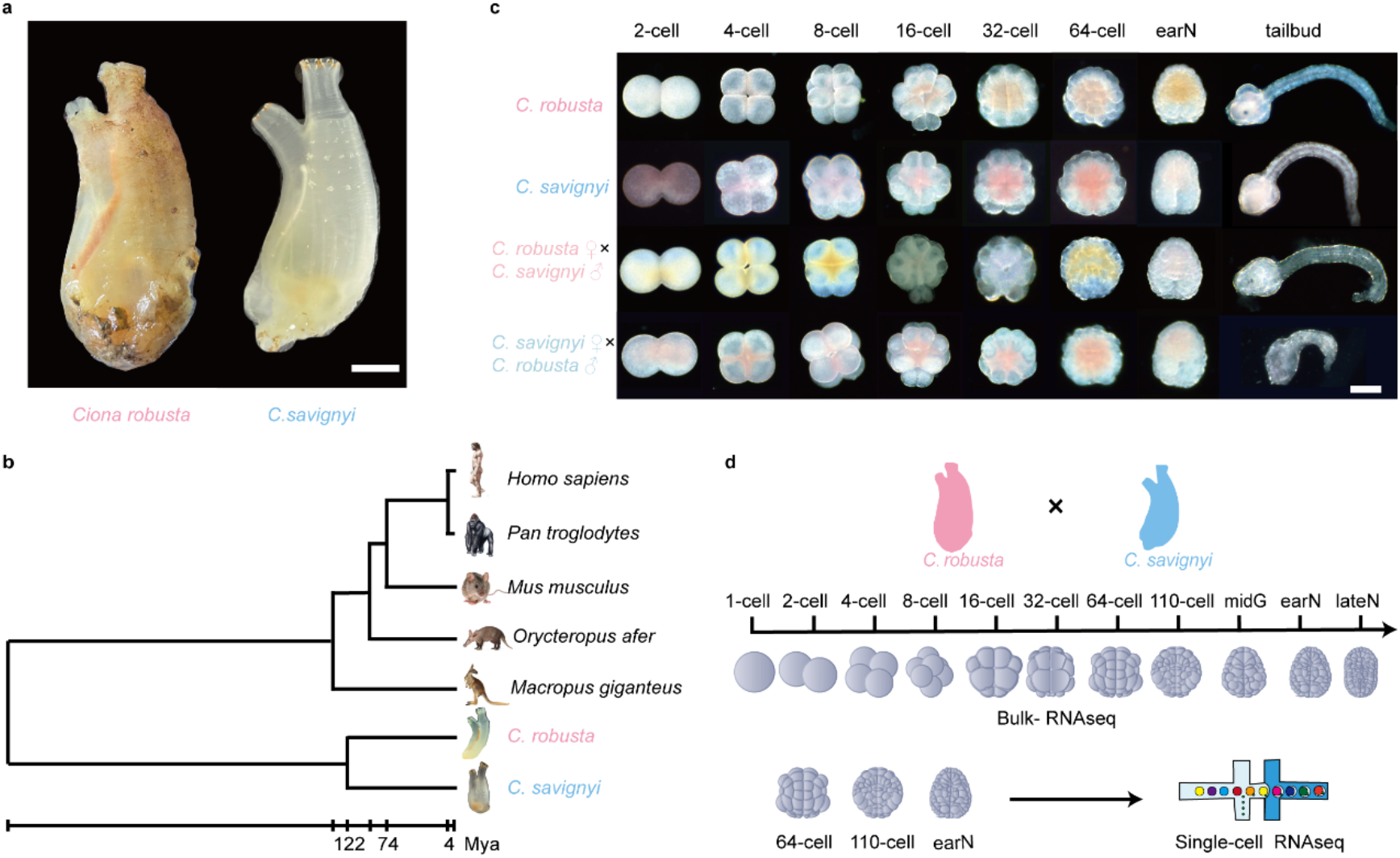
Generating hybrid and gene expression profiles from two divergent *Ciona* species. **a**, Morphology of adult *C. robusta* (left) and *C. savignyi* (right). Scale bar: 1 cm. **b**, Phylogenetic position and divergence time estimation of *Ciona*. Divergence times are displayed below the phylogenetic tree. **c**, Different developmental stages of four cross ascidians, including self-crossing *C. robusta*, self-crossing *C. savignyi, C. savignyi* as a male parent cross with *C. robusta* (Cs♂ × Cr♀), and *C. robusta* as a male parent cross with *C. savignyi* (Cr♂ × Cs♀). Eight developmental stages were tested, including 2-cell, 4-cell, 8-cell, 16-cell, 32-cell, 64-cell, early-neurula, and tailbud stages. Scale bar: 50 μm. **d**, Overview of the experimental strategy. Cross embryos of both Cs♂ × Cr♀ and Cr♂ × Cs♀ were collected for bulk RNA sequencing, whereas cross embryos at the 64-cell, 110-cell, and early-neurula stages were collected for single-cell RNA sequencing.

First, we assessed the developmental capabilities of the hybrid embryos based on the dechorionized eggs of the two species^22^. Embryos were obtained from each cross, and the embryogenesis of each cross involved a similar developmental time and morphology until the tailbud stage (Fig. 1c). Although one direction of the hybrid embryos (Cr♂ × Cs♀) had a higher development failure rate, there was no significant difference in the embryogenesis process between Cs♂ × Cr♀ and Cr♂ × Cs♀ embryos before the tailbud stage, which is distant stage from our research window for ZGA.

Although they are morphologically similar, the two tested species are highly divergent in terms of their genomic sequences. The identity of all aligned orthologous proteins is ∼85% (Extended Data Fig. 1), which allows us to distinguish two alleles from either male or female parents in hybrid experiments. Thus, to determine the spatial activation of zygotic transcripts, bidirectionally heterologous embryos at the 1-cell, 2-cell, 4-cell, 8-cell, 16-cell, 32-cell, 64-cell, 110-cell, mid-gastrula, early-neurula, and late-neurula stages were collected for bulk RNA-sequencing, and embryos at the 64-cell, 110-cell, and early-neurula stages were collected for single-cell RNA-sequencing (Fig. 1d).

Illumina short reads of bulk RNA-seq from each stage were mapped to the genomes of *C. robusta* and *C. savignyi* (Fig. 2a). For both Cs ♂ × Cr♀ and Cr ♂ × Cs♀ embryos, >99.99% of the mappable RNA-sequencing reads were mapped to the maternal genome in 1-cell-, 2-cell-, and 4-cell-stage sample, whereas paternal transcripts were detected from the 8-cell stage, and their proportion increased from the 16-cell to 32-cell stage (from 0.02% to 0.16% in Cs♂ × Cr♀ embryos and from 0.02% to 0.22% in Cr ♂ × Cs♀ embryo). In addition, the paternal reads ratio rose rapidly from the 64-cell to 110-cell stage (from 0.53% to 2.29% in Cs♂ × Cr♀ embryos and from 1.05% to 5.33% in Cr♂ × Cs♀ embryos; Fig. 2b). The disproportionate transcription of maternal and paternal genomes was diminished in the early-neurula and late-neurula stages. For example, at the late-neurula stage, 13.65% of reads were mapped to paternal genome in Cs♂ × Cr♀ embryos, whereas 26.90% of reads were mapped to paternal genomes in Cr♂ × Cs♀ embryos. However, the proportion of maternal reads remained higher than that of the paternal reads, indicating that maternally inherited transcripts might still be retained until the neurula stage.

**Fig. 2.**
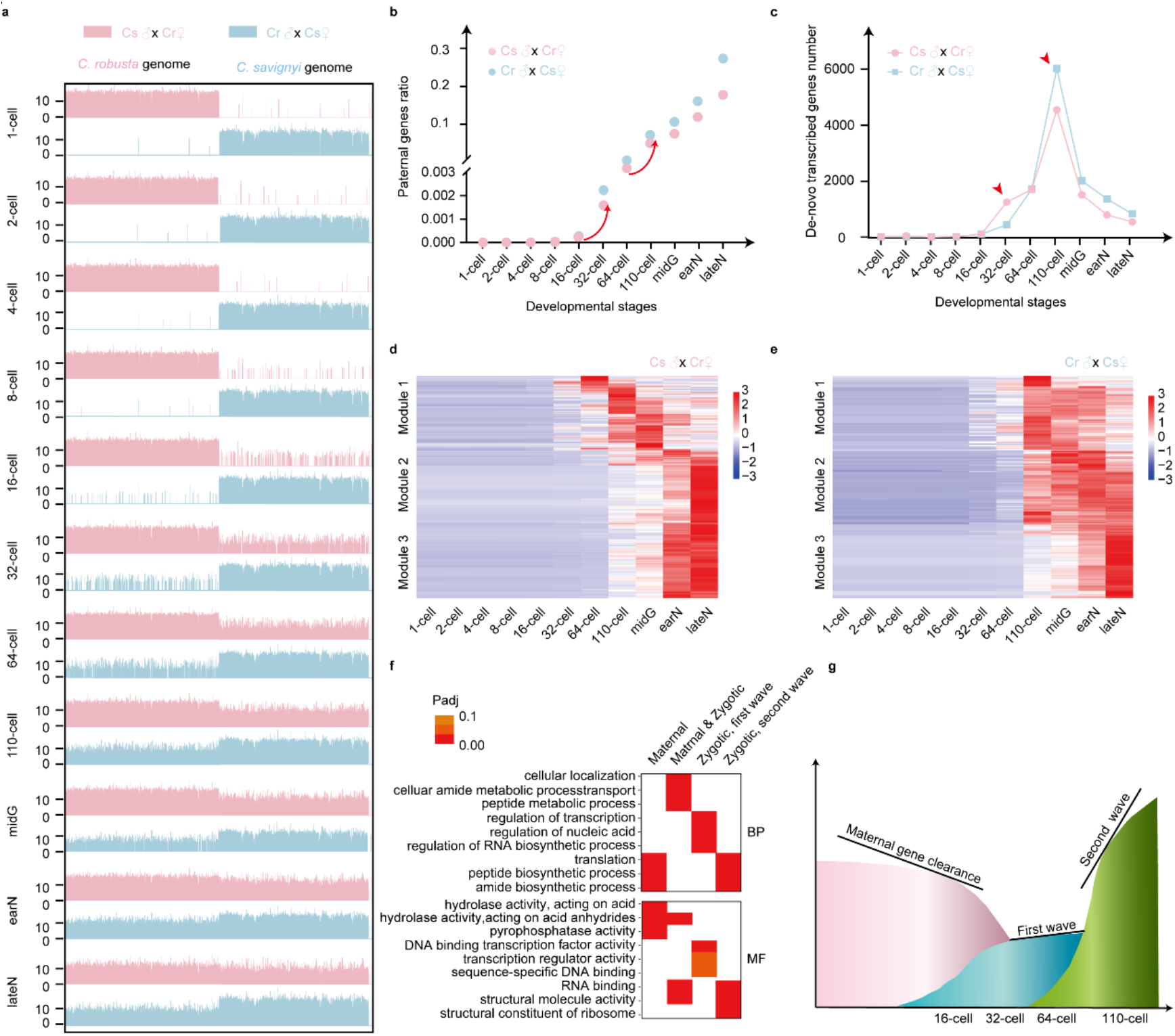
Two co-ordinated waves of ZGA during ascidian embryogenesis. **a**, Illustration of reads mapping to the genomes of *C. robusta* and *C. savignyi* at different developmental stages. Horizontal and vertical axes represent genome position and log_2_-transformed read number in each window, respectively. **b**, Gene expression ratios of paternal genes to all genes at different developmental stages. **c**, Number of *de-novo* transcribed genes at different developmental stages. **d**, Heatmap of paternal gene expression in different modules in Cs♂ × Cr♀ embryos. Module 1 included first wave genes, the expression of which increased from the 16-cell to 32-cell stage, whereas modules 2 and 3 included second wave genes, the expression of which increased from the 64-cell to 110-cell stage. **e**, Heatmap of paternal gene expression in different modules in Cr♂ × Cs♀ embryos. The description of the modules given in **d** also applies here. **f**, GO enrichment analysis. Left, biological process; right, molecular function. Color indicates adjusted p-values. **g**, Model of gene expression profiles during early developmental stages in *Ciona* embryogenesis. Details are provided in the Supplementary Table.

We measured gene expression levels using fragments per kilobase of genes per million mapped reads (FPKM) values, which were set at 0.1 as a cutoff for determining gene expression. The first expression of zygotic genes occurred at the 8-cell stage in both Cs♂ × Cr♀ and Cr♂ × Cs♀ embryos. These genes were *foxA2, regulator of G-protein signaling 8*, and *pbx1* in Cs♂ × Cr♀ embryos, and *foxA2, Sox2, SH3 domain-binding protein 4, DNA-directed RNA polymerase I subunit RPA1, brachyury*, and *regulator of G-protein signaling 8* in Cr♂ × Cs♀ embryos. Analysis of the occurrence stage of *de novo* transcribed paternal genes also showed an increase in states both from the 16-cell to 32-cell stage and from the 64-cell to 110-cell stage (Fig. 2c), consistent with a previous study^23^.

Principal component analysis divided the bulk RNA-sequencing samples into four quadrants (Extended Data Fig. 2). The distinction from the 64-cell to 110-cell stage was consistent with the paternal read ratio and *de novo* transcribed gene number analysis. Combining the appearance of gene and expression characteristics, the initiation of ZGA in *Ciona* began at the 8-cell stage, the first wave was from the 16-cell to 32-cell stage, and the second wave was from the 64-cell to 110-cell stage.

To ensure whether maternal orthologs were equally transcribed as paternal’s detected, we further performed CUT&Tag for RNA polymerase II (Pol II) and H3K27ac (a histone modification associated with active enhancer) in Cs♂ × Cr♀ embryos. The results showed that Pol II were equally recruited and elongated at parental orthologs for those early zygotic genes and reactivated housekeeping genes, along with H3K27ac deposited (Extended Data Fig. 3a, b). These CUT&Tag results support the idea that paternal transcripts could be used for studying housekeeping gene reactivation with little artifacts derived from cross.

**Fig. 3.**
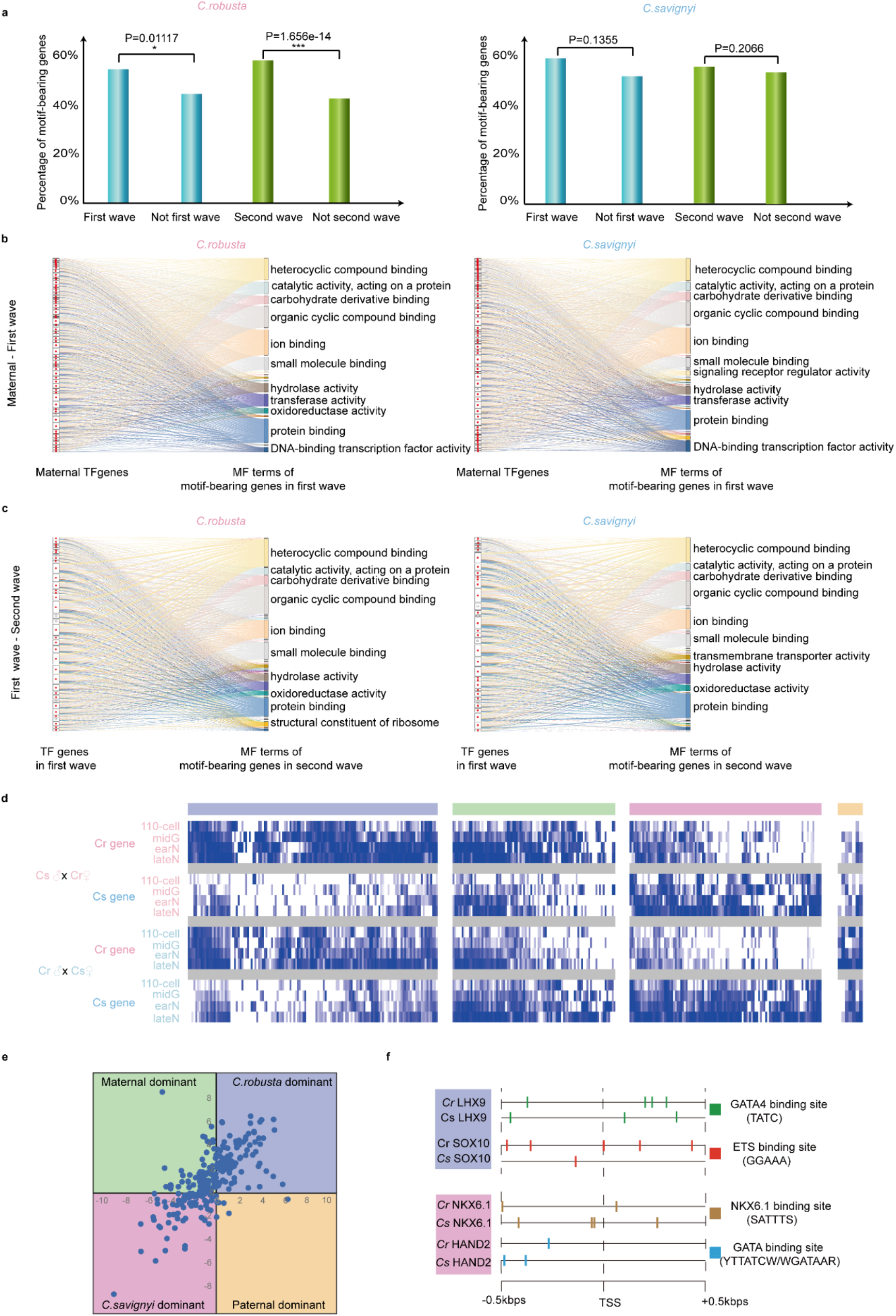
Allelic gene activation during embryogenesis of hybrid animals. **a**, Proportion of motif-bearing genes in the first and second waves. Asterisks represent significant difference with the control group. * *P* < 0.05 and *** *P* < 0.001 by Students *t*-test. **b**, Regulation relationships between maternal TFs and their motif-bearing genes in the first wave. Left side, maternal TFs; solid red dots show maternal TFs that are shared between *C. robusta* and *C. savignyi* whereas unfiled dots show maternal TFs that are not shared. Right side, GO terms of motif-bearing genes in the first wave. **c**, Regulation relationships between TFs in the first wave and their motif-bearing genes in the second wave. Left side, maternal TFs; solid red dots show TFs that are shared in the first wave between *C. robusta* and *C. savignyi*, whereas, unfiled dots show TFs that are not shared. Right side, GO terms of motif-bearing genes in the second wave. **d**, Expression heatmap of genes in different quadrants from the 110-cell to late-neurula stage. **e**, Gene expression preference quadrant from the 110-cell to late-neurula stage. Quadrant is divided into four parts corresponding to four colors. X axis indicates the expression ratio between genes from *C. robusta* and the allelic genes from *C. savignyi* in Cr♂ × Cs♀ embryos. Y axis indicates the expression ratio between genes from *C. robusta* and the allelic genes from *C. savignyi* in Cs♂ × Cr♀ embryos. Therefore, the first and third quadrants indicate gene expression that is *C. robusta*-dominant and *C. savignyi*-dominant, respectively, whereas the second and fourth quadrants indicate gene expression that is maternal-dominant and paternal-dominant, respectively. **f**, Comparison of motif distribution in *LHX9, SOX10, NKX6*.*1*, and *HAND2* between *C. robusta* and *C. savignyi*. Details are provided in the Supplementary Table.

Weighted correlation network analysis (WGCNA) of paternal genes showed that genes were clustered into five modules in Cs♂ × Cr♀ embryos (Extended Data Fig. 4a, b). The genes in module 1 were mostly upregulated from the 16-cell to 32-cell stage, whereas the genes in modules 2 and 3 were upregulated from the 64-cell to 110-cell stage (Fig. 2d and Extended Data Fig. 4c–f). A heatmap of paternal gene expression also showed that the module 1 contained genes in the first wave of ZGA, whereas modules 2 and 3 contained genes in the second major wave of ZGA (Fig. 2d). Gen Ontology (GO) enrichment analysis of each module showed that the genes in module 1 (first wave genes) were enriched in regulation, implying their regulatory effects on the second major wave genes (Fig. 2f). WGCNA and GO enrichment analysis of modules in Cr♂ × Cs♀ embryos showed similar results (Fig. 2e, f).

**Fig. 4.**
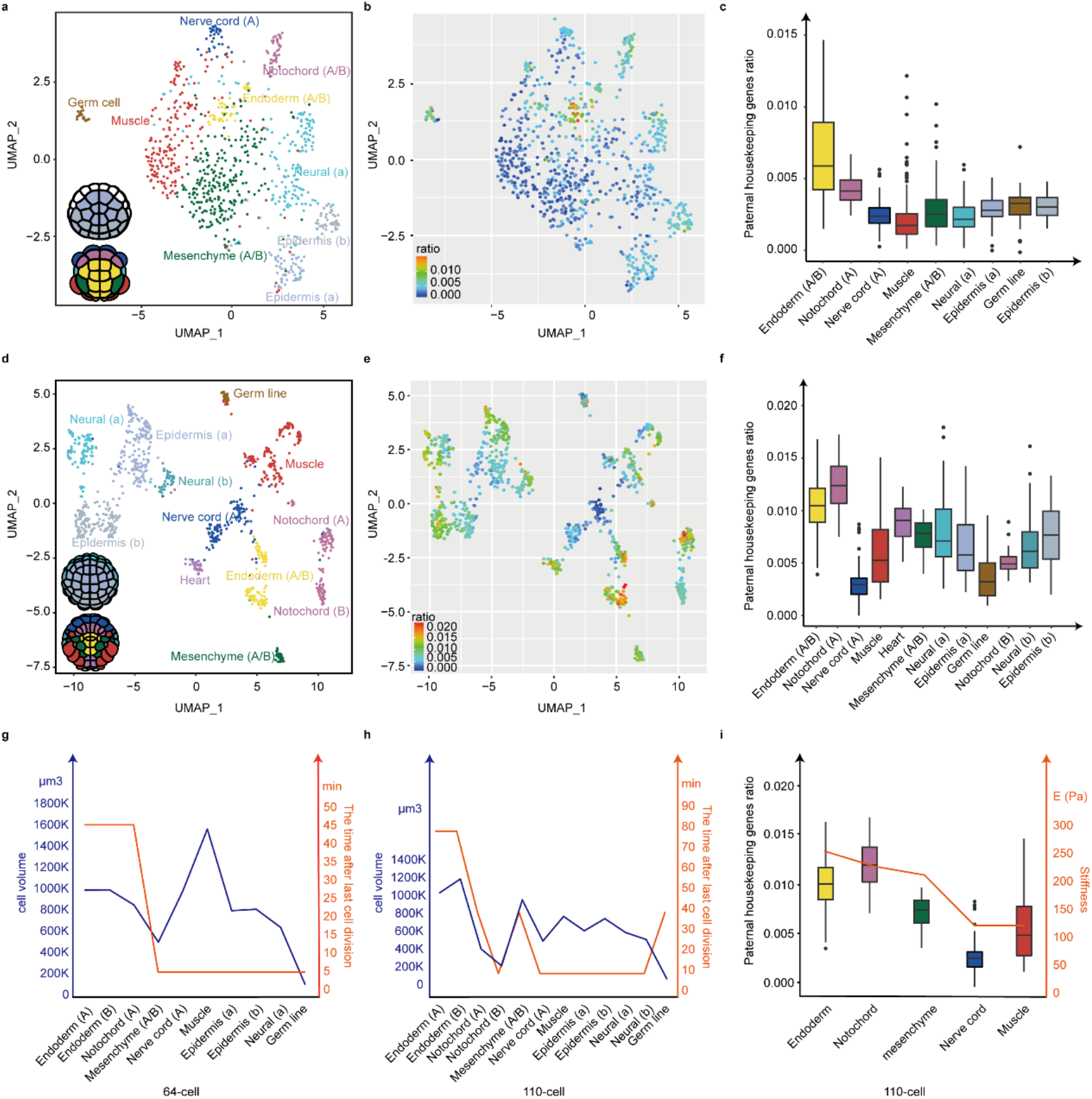
Housekeeping gene activation revealed by single-cell transcriptomics. **a**, UMAP plot of the 64-cell stage samples. In total, 844 cells were divided into 9 clusters and annotated as 8 cell types. The animal (up) and vegetable (down) blastomeres of a *Ciona* embryo at the 64-cell stage are shown. Font color corresponds to cell types. **b**, Expression ratio of paternal housekeeping genes in each cell at the 64-cell stage. Color represents the expression ratio. **c**, Boxplot of the expression ratio of paternal housekeeping genes in each cell type at the 64-cell stage. **d**, UMAP plot of the 110-cell-stage samples. In total, 1002 cells were divided into 12 clusters and annotated as 9 cell types. Animal (up) and vegetable (down) blastomeres of a *Ciona* embryo at the 110-cell stage are shown. Font color corresponds to cell type. **e**, Expression ratio of paternal housekeeping genes in each cell at the 110-cell stage. Color represents the expression ratio. **f**, Boxplot of the expression ratio of paternal housekeeping genes in each cell type at the 110-cell stage. **g**, Cell size and time after the last cell division in each cell type at the 64-cell stage. Blue and orange represent cell size and the time after the last time cell division, respectively. **h**, Cell size and time after the last cell division in each cell type at the 110-cell stage. Blue and orange represent the same variable as in **g. i**, Cell stiffness and boxplot of the expression ratio of paternal housekeeping genes in different cell types at the 110-cell stage. Orange line represents the stiffness values. Details are provided in the Supplementary Table.

Overall, the gene expression profiles of hybrid embryos indicated two co-ordinated waves of ZGA, especially from the 16-cell to 32-cell stage and from the 64-cell to 110-cell stage (Fig. 2g).

Next, we compared the regulatory relationships among the maternal, first wave, and second wave genes of *C. robusta* and *C. savignyi*. The general patterns of possible regulation relationships in the two species were similar (Fig. 3a–c; Extended data Figs. 5 and 6). First, we identified the transcription factor (TF) genes for which the corresponding motifs were well annotated, after which we traced the potential regulation between these TFs and genes that were potentially regulated, i.e., those with a motif located in their upstream sequence. Corresponding to maternal TFs, the proportion of motif-bearing genes in the first wave was higher than that in genes outside the first wave genes; corresponding to TFs in the first wave, the proportion of motif-bearing genes in the second wave was higher than that in genes outside the second wave genes (Fig. 3a). Motif analysis indicated that regulation of genes in the first wave was correlated with maternal TFs, whereas regulation of genes in the second wave was correlated with zygotic expressed genes in the first wave. The hybrid *Ciona* allowed us to compare the regulation patterns of developmental-related genes between *C. robusta* and *C. savignyi*. The relationship between maternal TFs and their motif-bearing genes in the first wave was similar between *C. robusta* and *C. savignyi*, i.e., maternal TFs were similar and the molecular functions, as well as biological processes and cellular components, of their downstream genes also fell into similar categories (Fig. 3b; Extended data Figs. 5a and 6a). Likewise, the relationship between TFs in the first wave and their motif-bearing genes in the second wave was similar (Fig. 3c; Extended data Fig. 5b and 6b), despite the long divergence time between the two species.

The sequence discrepancy between the two parental genomes also allows us to examine the expression preference between two alleles inherited from either male or female parents, i.e., allele-specific expression (ASE). Among 5,879 orthologous gene pairs detected, 341 pairs showed zygotic-only expression in both Cr♂ × Cs♀ and Cs♂ × Cr♀ embryos. We divided these pairs into four groups according to their expression ratio between *C. robusta* and *C. savignyi*. The early-expressed genes (from the 8-cell to 32-cell stage) (Extended Data Fig. 7a) tended to be *C. robusta*- or maternal-dominant, which might be attributed to the advantage of maternal inherited TFs at recognizing maternal enhancers. During the transition stage (the 64-cell and 110-cell stage) (Extended Data Fig. 7b), more *C. savignyi*- and paternal-dominant genes were detected. Of the late-expressed genes (from the 110-cell to late-neurula stage), 138 were *C. robusta*-dominant and 106 were *C. savignyi*-dominant in both Cr♂ × Cs♀ and Cs♂ × Cr♀ embryos, whereas 90 and 7 genes were maternal- and paternal-dominant, respectively, in both Cr♂ × Cs♀ and Cs♂ × Cr♀ embryos (Fig. 3d, e). These gene pairs showed the expression differences between allelic genes, especially from the 110-cell to late-neurula stage. Increased allelic gene activation during embryogenesis in hybrid animals tended to be influenced in a species-of-origin rather than parent-of-origin manner. We inferred that these species-of-origin effect allele-genes were related to variation in differential cis-regulatory elements. Thus, we compared the regulatory element binding motif in a 500-bp window flanking both sides of the transcription start site in 341 genes. A biased motif distributing pattern was identified in some ASE genes subject to species-of-origin effects. The distribution of the trans-regulating element binding motifs of *LHX9*^24^ and *SOX10*^25^ in homolog genes in *Ciona* showed a *C. robusta* dominant pattern, whereas motifs regulating *NKX6*.*1*^26^ and *HAND2*^27^ showed a *C. savignyi*-dominant pattern. These findings indicate that the biased expression of ASE may be attributable to the divergence of cis-regulatory elements divergence between allelic genes. GO enrichment analysis of ASE genes revealed that species-of-origin determinant genes were enriched in the immune system process and ion transport, whereas parent-of-origin determinant genes were enriched in the metabolic process and regulation of transcription process (Extended Data Fig. 8). These findings are consistent with previous studies, in which many ASE genes in humans were associated with the immune response^28^, while parent-of-origin effects were enriched on metabolic traits in mouse^29^. The allelic gene activation during embryogenesis of hybrid *Ciona* showed that 72% ASE genes were species-of-origin effects, the other 28% ASE genes were parent-of-origin effects. Evidence from large-scale studies in mammals also suggested that the majority of the imbalance of ASE resulted from genetic effects rather than imprinting or random monoallelic expression^30^.The asynchronized division of *Ciona* is initiated at the 16-cell stage, which is earlier than the initiation of housekeeping gene reactivation. The mosaic development with invariant lineage in *Ciona* embryos provides a unique opportunity to study ZGA and test two prevailing models simultaneously at a single cell resolution. Thus, we examined the reactivation of paternal housekeeping genes (modules 2 and 3) using single-cell RNA-sequencing of 64-cell to early neurula stage embryos, i.e., when the major ZGA wave occurred. Synchronized Cs♂ × Cr♀ embryos at the 64-cell and 110-cell stages were rapidly dissociated in ice-cold RNase-free calcium-free synthetic seawater and processed in a 10× Genomics Chromium system. In total, 8,357 single-cell transcriptomes were obtained after filtering. Cells from 64-cell, 110-cell, and early-neurula stage embryos were clustered into 9, 12, and 19 clusters using Uniform Manifold Approximation and Projection for Dimension Reduction (UMAP). Cell cluster identities were annotated as endoderm, notochord, mesenchyme, muscle, nerve cord, germ, neural, and epidermis (Fig. 4a, d; Extended Data Figs. 9–11) using published gene expression patterns^31, 32^.

To compare the ZGA in each cell cluster, we calculated the paternal transcript ratio of the housekeeping genes in each cell. The endoderm cells had the highest paternal transcript ratio, followed by notochord cells, then epidermis cells, muscle, nerve cord, and neural cells; mesenchyme and germline had the lowest ratio at the 64-cell stage (Fig. 4b, c; Extended Data Fig. 12a, b). At the 110-cell stage, endoderm cells and notochord cells had the top two highest paternal transcript ratios, whereas germ cells and nerve cord A lineage cells had the lowest ratios (Fig. 4e, f; Extended Data Fig. 12c, d). At the early-neurula stage, the paternal transcripts increased in nerve cord and mesenchyme B lineage cells, and the differences in the paternal transcript ratios of each lineage were narrowed (Extended Data Fig. 13). The differential reactivation of paternal housekeeping genes among different lineages suggests that the reactivation of housekeeping genes does not follow the developmental clock model (Fig. 4c, f).

We also analyzed the division number of each lineage to test the possibility of an N/C ratio model, in which the increasing cell cycle titrated maternally supplied repressors to relieve transcriptional repression. However, cell division time, which was indicated by the time after the last cell division in each cell lineage, was not correlated with the activation of zygotic genes (Fig. 4g, h). Nevertheless, it is possible that repressors existed as mRNA, and the rule of division may not have been followed exactly but might instead be reflected by cell size. Thus, we obtained cell volume data published previously^33^; However, we found that ZGA in hybrid *Ciona* was not correlated with cell volume (Fig. 4g, h). Therefore, neither the number of cell cycles nor the volume of lineages support the model of potential repressors. The area of contact between cells is also a key determinant of cell communication^33^. Given that endoderm invagination is the first event of gastrulation^34^, the earlier initiation of ZGA might be related to the cell movements and cell shape changes that follow. Previous studies have reported a large variation in cell mechanical properties at the single-cell level^35^. A study of the spatiotemporal dynamics of single-cell stiffness in the ascidian embryo revealed more stiffness in endoderm and adjacent notochord cells^36^. We compared the stiffness according to the expression of cell types and found that earlier ZGA in endoderm and adjacent notochord cells was indeed associated with stiffness properties in the early developing ascidian embryo (Fig. 4i). The expression profiles of stiffness signaling molecules, such as Rho, ephrin, and Nodal, were also correlated with ZGA in different cell lineages (Extended Data Fig. 14). In mammalian embryos, differential cell surface fluctuations reportedly account for the lineage sorting of the primitive endoderm from the epiblast^37^. The primitive endoderm displays higher surface fluctuations than the epiblast. These nonequilibrium cell surface dynamics could also be associated with the asynchronized ZGA in *Ciona*.

By taking advantage of the natural “genetic tags” from distinct *Ciona* hybrids, we revealed two waves of ZGA, alone and with housekeeping gene reactivation, from the 16-cell to 32-cell stage and from the 64-cell to 110-cell stage. Furthermore, the hybrid ascidians also provide a potentially powerful system to study the evolution of cis- and trans-regulation during embryogenesis. we found that species-of-origin rather than parent-of-origin ASE effects occurred during ascidian embryogenesis. Intriguingly, we identified a different ZGA pattern, in which the housekeeping gene reactivation could not be explained by either absolute timing, or the N/C ratio, showing a significant correlation with cell stiffness properties. These finding support the notion that ZGA, especially the reactivation of housekeeping genes, is under extreme selection pressure, and our study provides a new system for understanding the evolution and adaptation of strategies that regulate this critical developmental process.

## Methods

### Animal hybrids and sample collection

Adults *C. robusta* and *C. savignyi* were collected in Weihai, China and bred in sea water at 18°C in the laboratory. Eggs and sperm were obtained from mature animals as described previously^22^. We crossed two animals: *C. savigyni* sperm was mixed with *C. rubusta* eggs, with the resultant embryos defined as Cs♂ × Cr♀, and the *C. robusta* sperm was mixed with *C. savignyi* eggs, with the resultant embryos defined as Cr♂ × Cs♀. After 10 min, the fertilized eggs were washed twice with filtered sea water. The embryos were raised to different stages at 18 °C^38^. We collected Cs♂ × Cr♀ and Cr♂ × Cs♀ crossed embryos at the 2-cell, 4-cell, 8-cell, 16-cell, 32-cell, 64-cell, 110-cell, mid-gastrula, early-neurula, and late-neurula stages for morphological observation and sequencing.

### Bulk RNA-sequencing and data analysis

For each sample, 100–500 morphologically normal embryos were randomly selected and transferred into tubes precoated with 0.1% bovine serum albumin (BSA). Total RNA was extracted using TRIzol RNA extraction reagent. A Nanodrop spectrophotometer and Agilent 2100 bioanalyzer were used to assess RNA integrity. RNA samples were sent to Novogene company (China) for transcriptome sequencing via an Illumina NovaSeq system in paired-end 150-bp mode. Reads were aligned to the *C. robusta* (KY2019; Ghost, http://ghost.zool.kyoto-u.ac.jp/default.html) and *C. savignyi* (CSAV2.0; Ensembl, https://asia.ensembl.org/index.html) genomes using HISAT2 software v2.0.5^39^. Gene expression levels were estimated using featureCounts (1.5.0-p3)^40^ to acquire the FPKM value of each gene. A coexpression gene network for 22 transcriptomic datasets was constructed using the R package WGCNA^41^. Experimentally verified trans-acting factor binding sites or motifs of ascidians were acquired from JASPAR^42^. TBtools (v1.098696)^43^ was used to select these upstream sequences. The mast tool (v5.0.5)^44^ implemented in the meme suite^45^ was used to search the their upstream 1,000-bp sequences for matches to motifs.

### Bulk CUT&Tag and data analysis

200 uL Ca^2+^-free artificial sea water (ASW) containing 0.1% Triton X-100 was added into tubes containing embryos, which was centrifuged for 10 min at 1300 g. Nucleus were extracted from hybrid 64-cell stage embryos. CUT&Tag was performed on pre-washed nucleus with anti-RNA polymerase II antibody and anti-H3k27ac antibody as described^46^. All sequencing data were aligned to the *C. robusta* and *C. savignyi* genomes using Bowtie2 version 2.4.4^47^. Bam file produced by bowtie2 were transferred to browsable bw file with bamCoverage version 3.5.1 for next step analysis. Genome browser snapshot was captured with Integrative Genomics Viewer (IGV, v2.12.2)^48^ using Run Batch Script tool. DeepTools (v3.5.1)^49^ was used to plot peak distribution with computeMatrix scale-regions and plotHeatmap command.

### Cell dissociation of cross embryos

Morphologically normal embryos at the 64-cell, 110-cell, and early neural stage were collected and transferred into tubes precoated with 5% BSA in Ca_2_^+^-free ASW. Embryos were immediately dissociated using 0.2% trypsin in Ca_2_^+^-free ASW with 5 mM EGTA, after which they were pipetted for 3 min on ice to complete the dissociation of embryos into individual cells. Digestion was inhibited using 0.2% BSA in Ca^2+^-free ASW. Cells were collected via centrifugation at 4 °C and 500 g for 2–5 min and resuspended in ice-cold Ca^2+^-free ASW containing 0.1% BSA.

### Single-Cell RNA-Sequencing library construction and sequencing

For 64-cell- and 110-cell-stage embryos, single-cell gel beads in emulsion were generated using a Chromium Controller instrument (10× Genomics). Sequencing libraries were prepared using Chromium Single-Cell 3′ Reagent Kits (10× Genomics) according to the manufacturer’s instructions. After performing cleanup using a SPRIselect Reagent Kit, the libraries were constructed by performing the following steps: fragmentation, end-repair, A-tailing, SPRIselect cleanup, adaptor ligation, SPRIselect cleanup, sample index PCR, and SPRIselect size selection. For early neurula-stage embryos, sample preparation, library construction, and single-cell sequencing were performed using a BD Rhapsody Single-Cell Analysis, for which cell capture beads were prepared and loaded onto the cartridge. According to the manufacturer’s protocol, cartridges were washed, cells were lysed, and cell capture beads were retrieved and washed prior to performing reverse transcription and treatment with Exonuclease I. Subsequently, cDNA libraries were prepared using mRNA-targeted sample tags, and a BD AbSeq library was prepared using the BD Rhapsody-targeted mRNA and AbSeq Amplification Kits. Libraries were sequenced using an Illumina NovaSeq system.

### Single-cell RNA-sequencing data analysis

For 10× Genomics data, bcl files were converted into FASTQ format using bcl2fastq v1.8.4 (Illumina). To generate gene– barcode matrices for each sample, we used 10× Cell Ranger v3.0.2 with its default settings. For BD Rhapsody data, the Seven Bridges platforms pipe line v1.9 (https://www.sevenbridges.com) was used. We mapped sequencing reads to the reference genome combining both *C. robusta* and *C. savignyi*. To remove signals from hypothetical empty droplets or degraded RNA, a preliminary filter was performed to preserve all genes expressed in at least three cells as well as at least 250 genes expressed in a single-cell. We normalized the read counts of each cell using Seurat V4^50^, and 2000 highly variable genes were selected from each sample for downstream principal component analysis. The cells from 64-cell-stage embryos were filtered using 1000– 7000 detected genes, resulting in 844 cells with 3049 genes and 7892 unique molecular identifiers (UMIs) per cell (corresponding to 13.2-fold coverage). The cells from 110-cell-stage embryos were filtered using 2000–7000 detected genes, resulting in 1002 cells with 4836 genes and 19,012 UMIs per cell (corresponding to 9.1-fold coverage). The cells from early neurula-stage embryos were filtered using 500–4500 detected genes, resulting in 6511 cells with 2606 genes and 4888 UMIs per cell (corresponding to 21-fold coverage). Cell distance was visualized using the UMAP method in reduced two-dimensional space. We annotated the cell clusters according to the marker genes in each cell cluster. For 64-cell-stage samples, 9 cell clusters were annotated; for 110-cell-stage samples, 12 cell clusters were annotated; and for early neurula-stage samples, 19 clusters were annotated.

### Statistics analysis

All data statistics analysis was performed using paired Student’s t-tests. The P value < 0.05 was considered as significant difference exists. * represents 0.01 < P < 0.05. ** represents 0.001 < P < 0.01. *** represents P < 0.001.

## Data availability

All data generated or analyzed during this study are included in the manuscript and supporting files. The bulk RNA-sequencing and single-cell RNA-sequencing datasets generated in this study have been deposited at NCBI Gene Expression Omnibus under the accession numbers SRR18477895–SRR18477916 and SRR18494244– SRR18494246. All other data supporting the findings of this study are available from the corresponding authors on reasonable request. Source data are provided with this paper.

## Author contributions

K.C. and B.D. conceived and designed the experiments. Most experiments and data analysis were carried out by J.W. and W.Z. with help from Y.L., Y.G., Q.Z., L.Z., and H.Y.. J.W., L.L., and A. J. performed the data analysis. Initial draft was drafted by J. W. and W. Z.. Final version of the manuscript was prepared by B.D., K.C.. All authors proved the manuscript.

## Acknowledgements

This work was supported by the National Key Research and Development Program of China (2019YFE0190900, 2018YFD0900705), and the Taishan Scholar Program of Shandong Province, China (to B.D.).

